# Ancient Migrations - The first complete genome assembly, annotation and variants of the Zoroastrian-Parsi community of India

**DOI:** 10.1101/2021.02.17.431362

**Authors:** Naseer Pasha, Kashyap Krishnasamy, Naveenkumar Nagarajan, Seshank Mutya, Bhavika Mam, Kouser Sonnekhan, Chellappa Gopalakrishnan, Renuka Jain, Villoo Morawala-Patell

## Abstract

With the advent of Next Generation Sequencing, many population specific whole genome sequences published thus far, predominantly represent individuals of European ancestry. While sequencing efforts of underrepresented communities in genomes datasets, like the Yoruba West-African, Han Chinese, Tibetan, South Korean, Egyptian and Japanese have recently added to the public genomic repositories, a comprehensive understanding of human genomic diversity and discovery of trait-associated variants necessitates the need for additional population specific analysis. In this context, the genomics of the population from the Indian sub-continent, given its genetic heterogeneity needs further elucidation.

In this context, the endogamous Zoroastrian-Parsi community of India, offer an exceptional insight into a homogenous population that has culturally, socially, and genetically remained intact, for 13 centuries amidst the genomic, social and cultural Indian landscape, consequent to their migration from the ancient Persian plateau.

Notwithstanding longevity as a trait, this endangered community is highly susceptible to cancers, rare genetic disorders, and display a documented high incidence of neurodegenerative and autoimmune conditions. The community as a matter of cultural practice abstains from smoking.

Here, we describe the assembly and annotation of the genome of an adult female, Zoroastrian-Parsi individual sequenced at a high depth of 173X using a combination of short Illumina reads (160X) and long nanopore reads (13X). Using a combination of hybrid assemblers, we created a new, population-specific human reference genome, The Zoroastrian-Parsi Genome Reference Female, AGENOME-ZPGRF, contains 2,778,216,114 nucleotides as compared to 3,096,649,726 in GRCh38 constituting 93.235% of the total genomic fraction. Annotation identified 20833 genomic features, of which 14996 are almost identical to their counterparts on GRCh38 while 5837 genomic features were covered in partial. AGENOME-ZPGRF contained 5,426,310 variants of which the majority were SNP’s (4,291,601) and 960,867 SNPs were AGENOME-ZPGRF specific personal variants not listed in dbSNP.

We present, AGENOME-ZPGRF as a whole reference for any genetic studies involving Zoroastrian-Parsi individuals extending their application to identify disease relevant prognostic biomarkers and variants in global population genomics studies.

## Introduction

Recent technological advances in high-throughput genome sequencing have brought a steep decline in the cost of genetic information^1^, while increasing the predictive power and path to clinical translation of risk estimates for common variants found in genome wide association studies^2^. Most massively parallel sequencing approaches use simple alignment of short reads to a reference genome to study genomic variation. While the approach has been successful^3^, an exhaustive study of structural variants and SNPs at a high depth, coverage, and confidence is essential for translation to precision medicine. The increase in long read sequencing technologies, as part of the 3^rd^ generation genomic approaches have facilitated the assembly of large eukaryotic genomes in the last decade^4,5^. These advanced genomic platforms have given us powerful methods to generate long reads that compliment short accurate reads like those from Illumina sequencing chemistry to complete gaps and get better contiguity for the overall human genome.

Medical genetics has taken a leap forward in personalized medicine with the information of whole genome sequence for inheritable conditions, birth defects and chromosomal disorders^6^. Personalized genome assembly has shed light on the effects that non-genetic, disease-linked etiologies like methylation of CpG base pair islands have on gene availability for transcription^7,8^. With the advances in genomic sequencing, the importance of understanding genetic variability across extant human genomic diversity has become crucial^9^, especially since the current reference genome assembly GRCh38^10^ and variants cover only a sub-section of global population sub-types due to its mosaic nature of Caucasian and African genomic admixtures. Therefore, approaches focusing on understanding the minor, ethnic population groups, hitherto unrepresented in major genome variation studies, such as HapMap^11^, 1000 Genome Initiative^12^, Human Genome Diversity Project^13^ and disease specific variation studies like TCGA^14^ has become a prerogative in population genomics. This approach has been extended to sequence whole genome population references from Chinese^15^, Ashkenazi^16^, Korean^17^, Japanese^18^, Turkish^19^, Egyptian^20^, South Indian Asian-Indian^21,22^ and many more draft genomes in the recent years. The availability of reference genomes from multiple human populations greatly aids attempts to find genetic causes of traits that are over- or under-represented in those populations, including susceptibility to disease.

The Indian subcontinent is a hotspot of social, ethnic and genetic diversity with waves of migration to Southeast Asia through India^23^. The genetic landscape of this region is mainly constituted from Austro-Asiatic (AA), Indo-European (IE), Tibeto-Burman (TB), and Dravidian (DR) families with cultural and social frameworks that discourage and at times prohibit intermarriages between ethnic groups^24,25^. The extensive genetic diversity of India with genetically isolated subpopulations, makes it an ideal for population genomic studies to explore the disease–variants relationships.

The Zoroastrian-Parsi of India represent one such endogamous, genetically homogenous community. While the community members have a longer median life span, their present numbers in India are dwindling making a genomic study of the community critical for population genomics. This community in India trace their origins to migrations from the Persian plateau (∼847 AD), from Pars and Khorasan through the island of Hormuz to India where they settled as Parsis and practiced their faith, Zoroastrianism. The venerate Fire as the medium of worship and practice ostracism against smokers, therefore representing an important genomic biobank in understanding diseases associated with nicotine dependence. The community has a high prevalence of cardiovascular disorders, Autoimmune disorders like Rheumatoid Arthritis, Neurological/Neurodegenerative conditions like Parkinson’s Disease, Alzheimes Disease and different types of cancers.

Here, we describe the assembly and annotation of the genome of an adult female, Zoroastrian-Parsi individual sequenced at a high depth of 173X using a combination of short Illumina reads (160X) and long nanopore reads (13X). Using a combination of hybrid assemblers, we created a new, population-specific human reference genome. The Zoroastrian-Parsi reference genome, AGENOME-ZPGRF, contains 2,778,216,114 nucleotides as compared to 3,096,649,726 in GRCh38. Annotation identified 20674 genomic features, of which 15235 are > 99% identical to their counterparts on GRCh38, while the remaining genes were found to covered in partial. AGENOME-ZPGRF contained 5,426,310 variants of which the majority were SNPs (4,291,601) and 960,867 SNPs were AGENOME-ZPGRF specific not listed in dbSNP.

## Materials and Methods

### Sample collection and ethics statement

The donor is a healthy, non-smoking Parsi female volunteer (age: 65 y.o), invited to attend blood collection camps at the Zoroastrian center in the city of Bangalore, India under the auspices of The Avestagenome Project^®^. The adult female (>18 years) underwent height and weight measurements and answered an extensive questionnaire designed to capture her medical, dietary, and life history. The subject provided written informed consent for the collection of samples and subsequent analysis. All health-related data collected from the cohort questionnaire were secured in The Avestagenome Project^®^ database to ensure data privacy.

### Genomic DNA extraction

Genomic DNA from the buffy coat of peripheral blood was extracted using the Qiagen Whole Blood and Tissue Genomic DNA Extraction kit (cat. #69504). Extracted DNA samples were assessed for quality using the Agilent Tape Station and quantified using the Qubit™ dsDNA BR Assay kit (cat. #Q32850) with the Qubit 2.0® fluorometer (Life Technologies™). Purified DNA was subjected to both long-read (Nanopore GridION-X5 sequencer, Oxford Nanopore Technologies, Oxford, UK) and short-read (Illumina Technologies)

### Library preparation and sequencing on the Nanopore platform

Libraries of long reads from genomic DNA were generated using standard protocols from Oxford Nanopore Technology (ONT) using the SQK-LSK109 ligation sequencing kit. Briefly, 1.5 µg of high-molecular-weight genomic DNA was subjected to end repair using the NEBNext Ultra II End Repair kit (NEB, cat. #E7445) and purified using 1x AmPure beads (Beckman Coulter Life Sciences, cat. #A63880). Sequencing adaptors were ligated using NEB Quick T4 DNA ligase (cat. #M0202S) and purified using 0.6x AmPure beads. The final libraries were eluted in 15 µl of elution buffer. Sequencing was performed on a GridION X5 sequencer (Oxford Nanopore Technologies, Oxford, UK) using a SpotON R9.4 flow cell (FLO-MIN106) in a 48-hr sequencing protocol. Nanopore raw reads (fast5 format) were base called (fastq5 format) using Guppy v2.3.4 software. Samples were run on two flow cells and generated a dataset of ∼14 GB.

### Library preparation and sequencing on the Illumina platform

Genomic DNA samples were quantified using the Qubit fluorometer. For each sample, 100 ng of DNA was fragmented to an average size of 350 bp by ultrasonication (Covaris ME220 ultrasonicator). DNA sequencing libraries were prepared using dual-index adapters with the TruSeq Nano DNA Library Prep kit (Illumina) as per the manufacturer’s protocol. The amplified libraries were checked on a Tape Station (Agilent Technologies) and quantified by real-time PCR using the KAPA Library Quantification kit (Roche) with the QuantStudio-7flex Real-Time PCR system (Thermo). Equimolar pools of sequencing libraries were sequenced using S4 flow cells in a Novaseq 6000 sequencer (Illumina) to generate 2 x 150-bp sequencing reads for 30x genome coverage per sample.

### Raw fastq files Illumina and nanopore reads

The sample genome was of an adult female from the endogamous Parsi community which was used for the construction of 173X Zoroastrian Parsi Whole Genome Assembly. Illumina HiSeq with a read length of 2 × 150 bp is used for obtaining short reads for the genome. We obtained a total of 2.2 Billion sequences from the Illumina HiSeq platform (160X) and a total of 6.8 Million reads from the Nanopore platform. For the long reads, the library preparation was according to the standard protocol and the sequencing of the genome was performed using the Oxford Nanopore Minion platform

### Quality trimming and Quality control of the reads

Quality trimming and adapter removal of the short Illumina platform reads was performed using AdapterRemoval (version 2.2.2)^26^ with minlength 30, trimwindow size 30 and reads lesser than quality score of Q30 were discarded. For adapter removal of long Oxford nanopore reads, Porechop tool (V0.2.4)^27^ with default options was used. The long error prone reads from Oxford Nanopore cannot cross the quality score of Q20 hence the cutoff was kept to 8 in this case. All the quality scores are checked using FastQC (version 0.11.5)^28^ and FastP (V0.20.1)^29^ tool (**Supplementary Figure 1, 2**).

### Whole Genome assembly

The quality trimmed and adapter removed short and long reads were processed for Hybrid assembly. The choice of hybrid assembly was made using relevant literature study where short read alone, long read alone and short-long read hybrid assemblies were compared for different cases^30^ and hybrid assemblies outperformed and gave better QC statistics reflecting better quality of the assembled genomes. The raw data was sub sampled to 60X coverage with length cutoff of 60 bp using fastP according to the instruction on the Wengan GitHub repository. The processed reads were assembled with Wengan^31^ using D mode (uses DiscovarDenovo short-read assembler) with options -l ontraw, -g 3000 (3Gbp). An alternative assembly was generated by using HASLR, Wtdbg2^32^, WenganA and WenganM assemblers.

### Removing mis-assemblies at segmental duplications and centromere regions

Centromere regions were downloaded from UCSC web browser for GRCh38 version of human reference genome. Segmental duplications in a BED format flat file was downloaded from the GitHub repository (segDupPlots/ucsc.collapsed.sorted.segdups). A python script20 was used to remove miss-assemblies from Segmental duplications and centromere regions. Identification and annotations of repetitive elements was obtained using REPEATMASKER (V4.1.1)33 by aligning the genome sequences against known library of repeats in humans.

### Read mapping and variant calling for Illumina sequencing reads

The variant detection for the AGENOME-ZPGRF female sample was carried out using GATK pipeline (V4.1.5.0)^35^, Picard (2.21.9) and Samtools (1.3.1)^36^. The GATK pipeline included read mapping and variant processing. Single-nucleotide variants (SNVs) and indels were called by local reassembly of haplotypes using HaplotypeCaller of GATK V4.1.5.0.

The following workflow was used for variant calling, the raw reads were pre-processed, converted to unaligned BAM and readgroup information were assigned using FastqToSam. The adapters were tagged using Markilluminadapter function and the bam file were converted to interleaved fastq sequences to map to the reference genome. The reference genome (GRCh38) is indexed using BWA index and samtools, further the sequence dictionary for reference was obtained using picard CreateSequenceDictionary function. Mapping the FASTQ reads to reference genome was performed by BWA-MEM (version 0.7.17-r1188). Information from unaligned BAM and the aligned BAM were merged using MergeBaMAlignment to retain the raw read information. The duplicate reads through experimental artefacts are tagged using MarkDuplicates (picard) module. The base quality score recalibration (BQSR) was applied to overcome the errors associated with base quality score due to sequencing errors. The BAM file were further indexed to identify variants by HaplotypeCaller. The variants obtained from HaplotypeCaller were annotated using SnpEff (4.3t), a genetic variant annotation and effect prediction toolbox.

### Structural variants

SVs were called using DELLY2^33^ with default parameters on duplicate marked bam file for germline SV calling (https://github.com/dellytools/delly).

### Pharmacogenomics relevance

To assess the pharmacogenomics relevance, we obtained common variants in AGENOME-ZPGRF and dbSNP-138 database. These variants were annotated based on PharmGKB (www.pharmgkb.org) database^34^ to obtain pharmacogenomics association. The variants that were classified as conflicting-interpretation, uncertain significance and benign were removed. The variants that had Pharmacokinetic (PK) and Phamacodynamic (PD) associations were considered to obtain actionable SNPs.

## Results

### Benchmarking of hybrid assemblers for Whole Genome Assembly using Chr22

*De novo* assembly was performed on a female Parsi whole-genome sequencing data. The data was in the form of Illumina-paired end 160X coverage and Oxford nanopore data of 13X. To standardize the pipeline, Chromosome 22 data (genome size of 52 Mbp) was extracted from the whole genome data and tested with iterative combination of different assembler strategy. The Illumina paired-end short reads with a read length of 150 bp were assembled using the Abyss^35^ assembler resulting in a total length of 44,331,422 bp and a contig N50 of 15,753 bp.

We proceeded to use Hybrid assemblers known to perform scaffolding using long reads and polishing using short reads. Wengan assembler outperformed all other hybrid assemblers by producing a total length of 32,346,746 bp with 194 contigs greater than 50,000 bp. Quickmerge^36^ meta-assembler was applied on Wengan assembly which gave the lowest number of contigs versus length and Abyss (160X) gave the longest length with the higher number of contigs improving the contiguity of the assembly. The results for this exercise produced a total length of 50,216,737 bp which is close to 90% of the total length of Chr22 and 2,675 contigs (**Appendix 1**).

### *De novo* assembly of the First complete Zoroastrian Parsi Whole Genome Reference Female (AGENOME-ZPGRF)

Following the assembly of Chr22, we extended our meta-assembly strategy to assemble the whole genome reference (**Figure 1**). Our Zoroastrian-Parsi genome (AGENOME-ZPGRF) is based on high-quality *de novo* assembly from one female Parsi individual. The assembly was generated from a combination of short and long read data sets: 2×150 bp Illumina paired-end reads (160X), Oxford Nanopore (13X) reads averaging over 5,784 bp in length (**Table 1**).

**Table 1:** Sequence data for assembly of *De novo* Zoroastrian-Parsi Genome Reference Female genome (AGENOME-ZPGRF)

| Sequence technology | Total reads (bp) | Mean read length(bp) | Coverage |
| --- | --- | --- | --- |
| Illumina-library-1 | 1422887753 | 151 | 100X |
| Illumina-library-2 | 322537951 | 150 | 30X |
| Illumina-library-3 | 479650511 | 151 | 30X |
| ONT-library-1 | 3318994 | 4566 | 5X |
| ONT-library-2 | 3515835 | 7002 | 8X |

**Figure 1:**
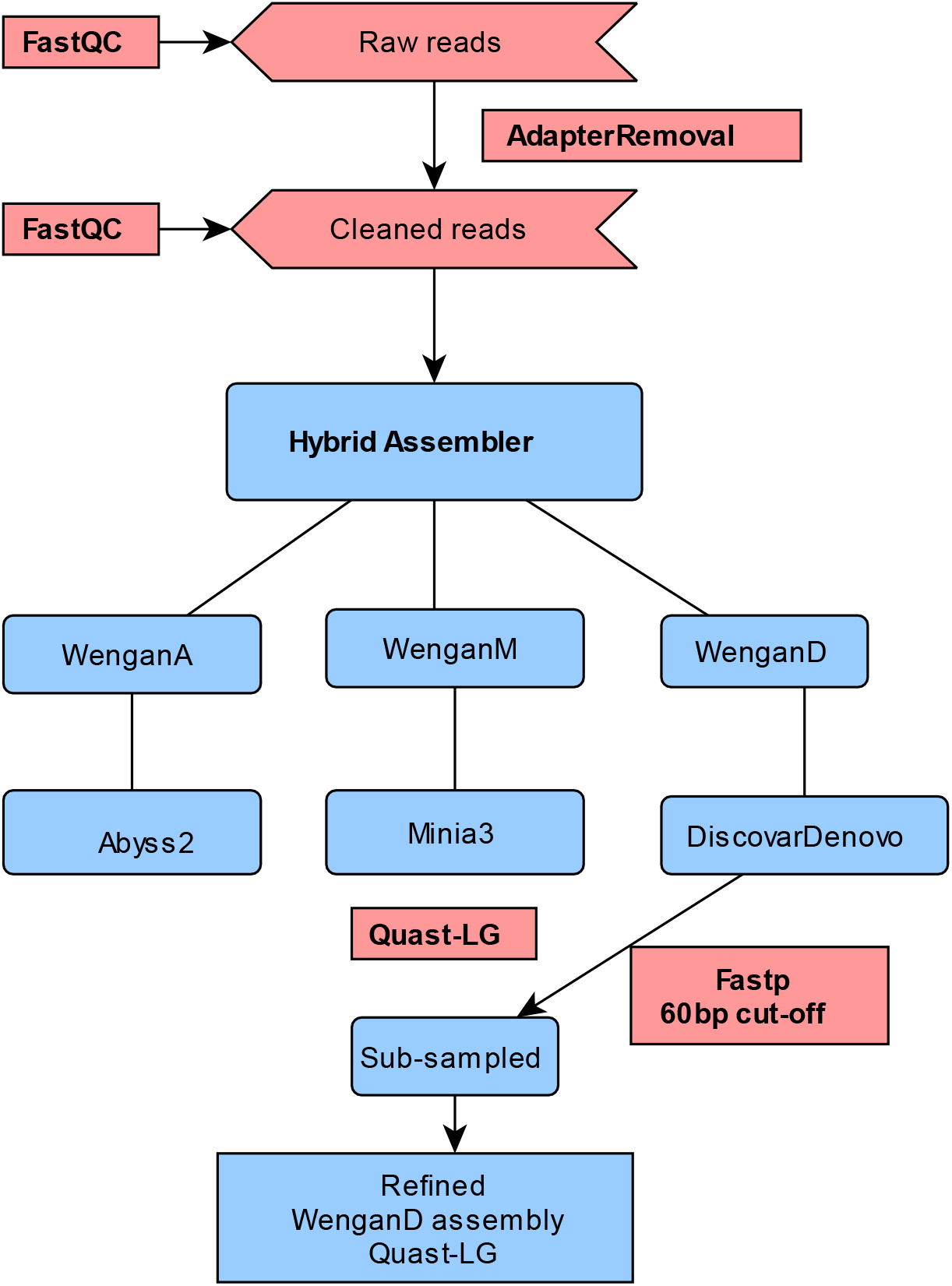
Workflow detailing meta-assembly protocol using iterative combinations of hybrid assemblers to generate the first Zoroastrian-Parsi Genome, AGENOME-ZPGRF

We initially created five hybrid assemblies using Illumina short reads and Oxford Nanopore Technology long reads. Three assemblies were based on Synthetic Scaffolding Graph approach using Wengan hybrid genome assembler, one assembly based on synthetic paired end reads using HASLR and one assembly based on fuzzy De-Brujin graph method using wtdbg2. WenganD gave the best results with respect to total length of 2.7 Giga bases (Gbp), N50 of 2 Mega bases (Mbp) and genomic fraction of 93.2%. The annotation based on QUAST-LG identified 14,996 complete and 5,837 partial number of genomic features (**Table 2**). The Parsi reference genome, AGENOME-ZPGRF, contains 2,778,216,114 nucleotides as compared to 3,096,649,726 in GRCh38. This assembly, designated AGENOME-ZPGRF, was the basis for all subsequent refinements and analysis.

**Table 2:** AGENOME-ZPGRF assembly statistics using Quast

| Genome statistics | AGENOME-ZPGRF | EGYPT | AK1 | YORUBA |
| --- | --- | --- | --- | --- |
| # contigs | 4806 | 3235 | 2832 | 1647 |
| # contigs (>= 0 bp) | 4806 | 3724 | 2832 | 1741 |
| # contigs (>= 1000 bp) | 4806 | 3560 | 2832 | 1726 |
| # contigs (>= 5000 bp) | 4775 | 2776 | 2832 | 1562 |
| # contigs (>= 10000 bp) | 4017 | 1917 | 2832 | 1347 |
| # contigs (>= 25000 bp) | 3335 | 1069 | 1570 | 799 |
| # contigs (>= 50000 bp) | 2966 | 734 | 747 | 288 |
| Largest contig | 15222969 | 88566048 | 113921103 | 248986603 |
| **Total length** | **2778216114** | **2836714529** | **2904207228** | **3088335497** |
| Total length (>= 0 bp) | 2778216114 | 2837486204 | 2904207228 | 3088495238 |
| Total length (>= 1000 bp) | 2778216114 | 2837367164 | 2904207228 | 3088485407 |
| Total length (>= 5000 bp) | 2778073401 | 2834831880 | 2904207228 | 3087990629 |
| Total length (>= 10000 bp) | 2772661904 | 2828723737 | 2904207228 | 3086359078 |
| Total length (>= 25000 bp) | 2761939683 | 2815431970 | 2882817238 | 3076876085 |
| Total length (>= 50000 bp) | 2748699427 | 2803817652 | 2855011855 | 3059626724 |
| **N50** | **2012108** | **25502944** | **44846623** | **155338310** |
| N75 | 937275 | 8329420 | 19924750 | 114367800 |
| L50 | 369 | 29 | 21 | 8 |
| L75 | 875 | 77 | 46 | 14 |
| GC (%) | 40.85 | 40.81 | 40.88 | 40.88 |
| **Misassemblies** |  |  |  |  |
| # misassemblies | 3322 | 1276 | 1952 | 1756 |
| # misassembled contigs | 777 | 484 | 782 | 374 |
| Misassembled contigs leng | 1398700724 | 2137050584 | 2657569650 | 3053643982 |
| # local misassemblies | 10110 | 10797 | 5004 | 5679 |
| # scaffold gap ext. mis. | 0 | 0 | 15 | 3 |
| # scaffold gap loc. mis. | 0 | 0 | 108 | 435 |
| # possible TEs | 250 | 330 | 256 | 296 |
| # unaligned mis. contigs | 33 | 333 | 455 | 228 |
| **Unaligned** |  |  |  |  |
| # fully unaligned contigs | 128 + 2501 part | 1207 | 402 | 412 |
| Fully unaligned length | 27948040 | 12541896 | 9234968 | 10329457 |
| **Genome fraction (%)** | **93.235** | **94.174** | **95.177** | **95.391** |
| Duplication ratio | 1.001 | 1.01 | 1.023 | 1.088 |
| **# genomic features** | **14996 + 5837 part** | **17682 + 3226 part** | **19651 + 1396 part** | **19356 + 1721 part** |
| Largest alignment | 9191143 | 75492126 | 58219133 | 65512502 |
| Total aligned length | 2749835393 | 2800100449 | 2829006639 | 2832740986 |
| NG50 | 1734895 | 20857787 | 39609866 | 145208384 |
| NG75 | 610541 | 5007910 | 14897232 | 114367800 |
| NA50 | 1339017 | 12942852 | 15098581 | 19529238 |
| NA75 | 659297 | 5094247 | 5183319 | 9114594 |
| NGA50 | 1175915 | 11187777 | 13028687 | 19529238 |
| NGA75 | 435352 | 3192669 | 3932304 | 8890200 |
| LG50 | 455 | 35 | 24 | 9 |
| LG75 | 1193 | 109 | 54 | 14 |
| LA50 | 584 | 60 | 59 | 43 |
| LA75 | 1316 | 146 | 140 | 96 |
| LGA50 | 712 | 71 | 66 | 43 |

### Completeness of the genome

Gene completeness was measured with BUSCO54 v.4.1.4^37^ using the Primates ODB-10 gene set. BUSCO provides intuitive metrics to describe genome, gene set or transcriptome completeness^38^. We observed 88.3% of genome completeness, 12,165 complete BUSCOs, 12,113 complete and single copy BUSCOs, 52 complete and duplicated BUSCOs, 461 fragmented BUSCOs, 1,154 missing BUSCOs. The total BUSCO group searched was 13,780.

### Repeated elements in AGENOME-ZPGRF

When annotating repeats with REPEATMASKER, about 48.34% of the genome (**Table 3**) was identified as repetitive, with its results similar to those from EGYPTRef, AK1 and YORUBA genome assemblies. Most of the repetitive elements comprised of 21.80% Long Interspersed Nuclear Elements (LINEs), 13.45% of Short Interspersed Nuclear Elements (SINEs) and the rest in ALU elements, Mammalian-wide Interspersed Repeats (MIRs), Long Terminal Repeats (LTR) elements, DNA elements, small RNAs, satellites and simple repeats. Further, we found that about two-thirds of the SNPs identified in the repeat regions were found in long interspersed elements (LINE; 21.80%; majority occurring in LINE1 elements) or short interspersed elements.

**Table 3:** Repetitive elements in AGENOME-ZPGRF identified using REPEATMASKER

| Repetitive elements | Number of elements | Length occupied | Percentage of sequence |
| --- | --- | --- | --- |
| **SINEs:** | 1592285 | 367249056 bp | 13.45 % |
| **ALUs** | 1078242 | 289089690 bp | 10.58 % |
| **MIRs** | 506412 | 77200621 bp | 2.83 % |
| **LINEs:** | 924026 | 595272136 bp | 21.80 % |
| LINE1 | 517835 | 482717596 bp | 17.67 % |
| LINE2 | 345948 | 98455152 bp | 3.60 % |
| L3/CR1 | 45096 | 10230478 bp | 0.37 % |
| **LTR elements:** | 485651 | 250839692 bp | 9.18 % |
| ERVL | 107081 | 55227511 bp | 2.02 % |
| ERVL-MaLRs | 241322 | 104424519 bp | 3.82 % |
| ERV_class I | 104268 | 76806564 bp | 2.81 % |
| ERV_class II | 6733 | 7378587 bp | 0.27 % |
| **DNA elements:** | 419798 | 102266826 bp | 3.74 % |
| hAT-Charlie | 214680 | 44096448 bp | 1.61 % |
| TcMar-Tigger | 96328 | 35713313 bp | 1.31 % |
| Unclassified: | 9436 | 4642192 bp | 0.17 % |
| **Total interspersed repeats:** |  | 1320269902 bp | 48.34 % |
| **Small RNA:** | 11816 | 1234995 bp | 0.05 % |
| **Satellites:** | 4807 | 10740343 bp | 0.39 % |
| **Simple repeats:** | 621517 | 35759325 bp | 1.31 % |
| **Low complexity:** | 94569 | 5807986 bp | 0.21 % |

### Variant identification

We used the GATK variant calling pipeline, performed according to GATK best-practice recommendations and the HaplotypeCaller tool was employed for identifying putative variants, followed by Snpeff (build 2017-11-24) to annotate and make functional predictions. Our analysis revealed 5,426,310 variants of which 79% are SNPs (4,291,601) and 21% are indels, multiple-nucleotide polymorphism (MNPs) and mixed variants (**Figure 2**). The transitions (297,330), transversions (251,540), Ts/Tv, ratio was 1.18, resembling expected figures in similar studies. Among the identified SNPs, 41.40% were intronic and 40.78% were intergenic while the rest were SNPs in the upstream (8.84%) and downstream region (7.19%). SNPs in the exonic region constituted only 0.5% of the total SNP count and the SNPs in the untranslated 3′ or 5′ region made up the rest (**Figure 2, Supplementary Figure 3**). Of these SNPs, 14,572 were missense (non-synonymous), 13,321 were silent (synonymous) substitutions and 189 nonsense SNPs. This is consistent with a non-syn:syn (dN/dS) ratio of ∼1 expected of a normal genome^39^.

**Figure 2:**
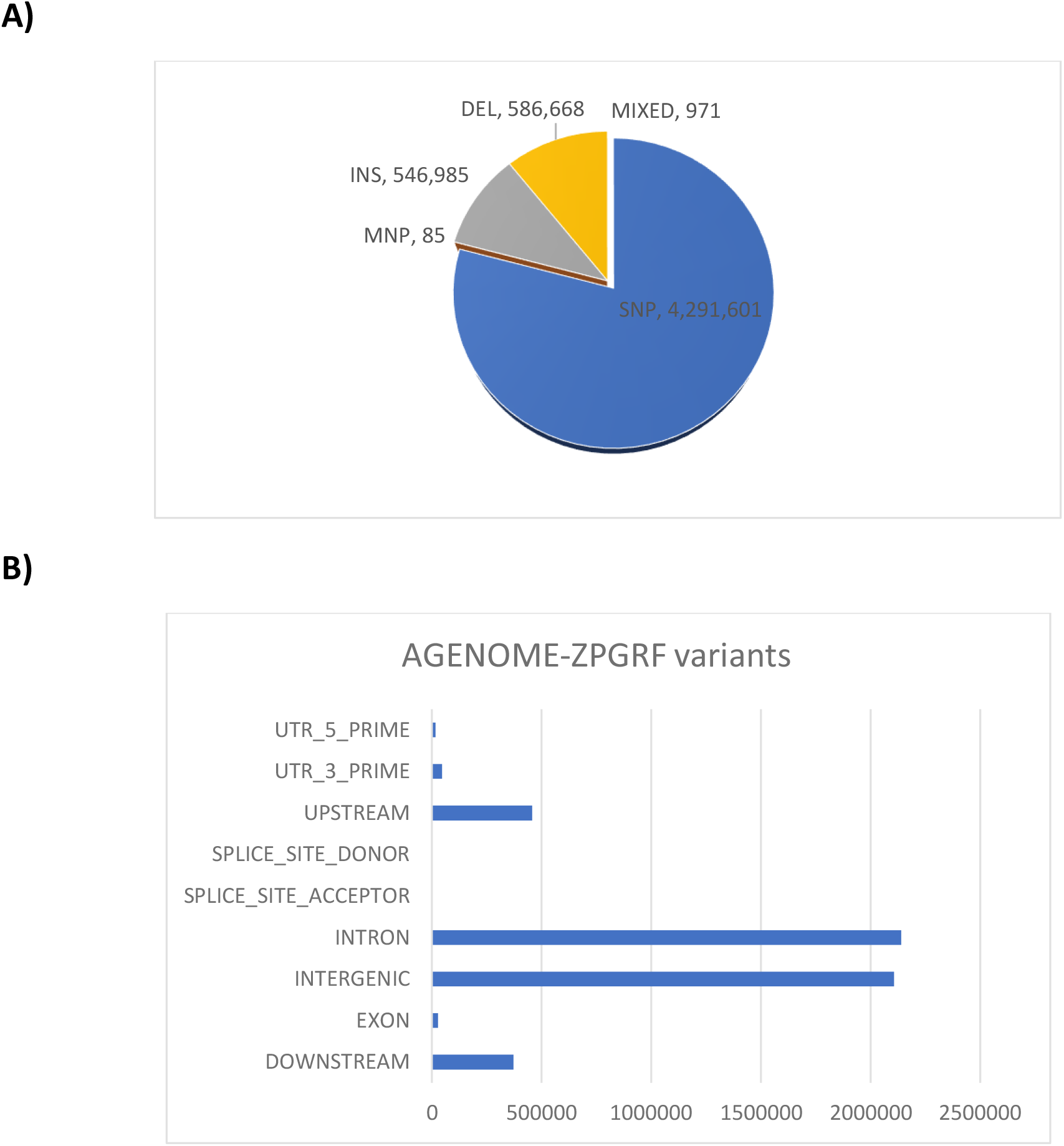
Distribution of variant types (A) and location in genomic regions (B) identified in the AGENOME-ZPGRF

Based on the SNPEff^40^ annotation, we sought to identify the functional impact of the SNP’s in terms of “high”, “low”, “moderate” impact based on their occurrence on the genome. High impact SNPs occur when (i) the variant hits a splice acceptor/donor site, (ii) a start codon is changed into a nonstart codon, or (iii) a stop codon is gained or lost due to the variant. We identified 1652 SNPs with a high-impact effect (**Appendix 2**), 16128 with low impact and 14856 with moderate impact. The majority of the high impact variants occurred on the gene *DPP6* (n=2920, **Appendix 4, 6**), variants of which have been reported to be associated to familial idiopathic ventricular fibrillation^41^.

Out of the ∼4.2 million SNPs identified, there were 960,867 potentially novel SNPs that did not exist in dbSNP^42^ (**Appendix 3, 7**). Further analysis of the AGENOME-ZPGRF specific novel SNPs showed, 50.9% and 31.2% were in intergenic and intronic regions, respectively. We found 9.2% were upstream, 7.17% downstream of a gene, and 4202 (or 0.49%) of the SNPs were found to be in coding regions (**Table 5**). Among the 4202 SNPs in coding regions, we could further classify 31 nonsense SNPs and a total of 415 SNPs with a high impact (**Appendix 5**). Most of the variants (both genomewide and AGENOME-ZPGRF specific variants) occurred on Chr1, while the highest frequency of distribution occurred on Chr22 (**Table 4**). We identified 1,133,653 indels, which consisted of 546,985 insertions and 586,668 deletions. Of these indels, 503,214 (or 44%) were found to be novel. Majority of the unique variants occurred on LOC105379427 (n=1552, **Appendix 4**) that code for zinc finger protein 717-like proteins (putative) and *DUX4L18/DUX4L19/CNTNAP3B* genes which have been implicated in cognitive disorders.

**Table 4:**
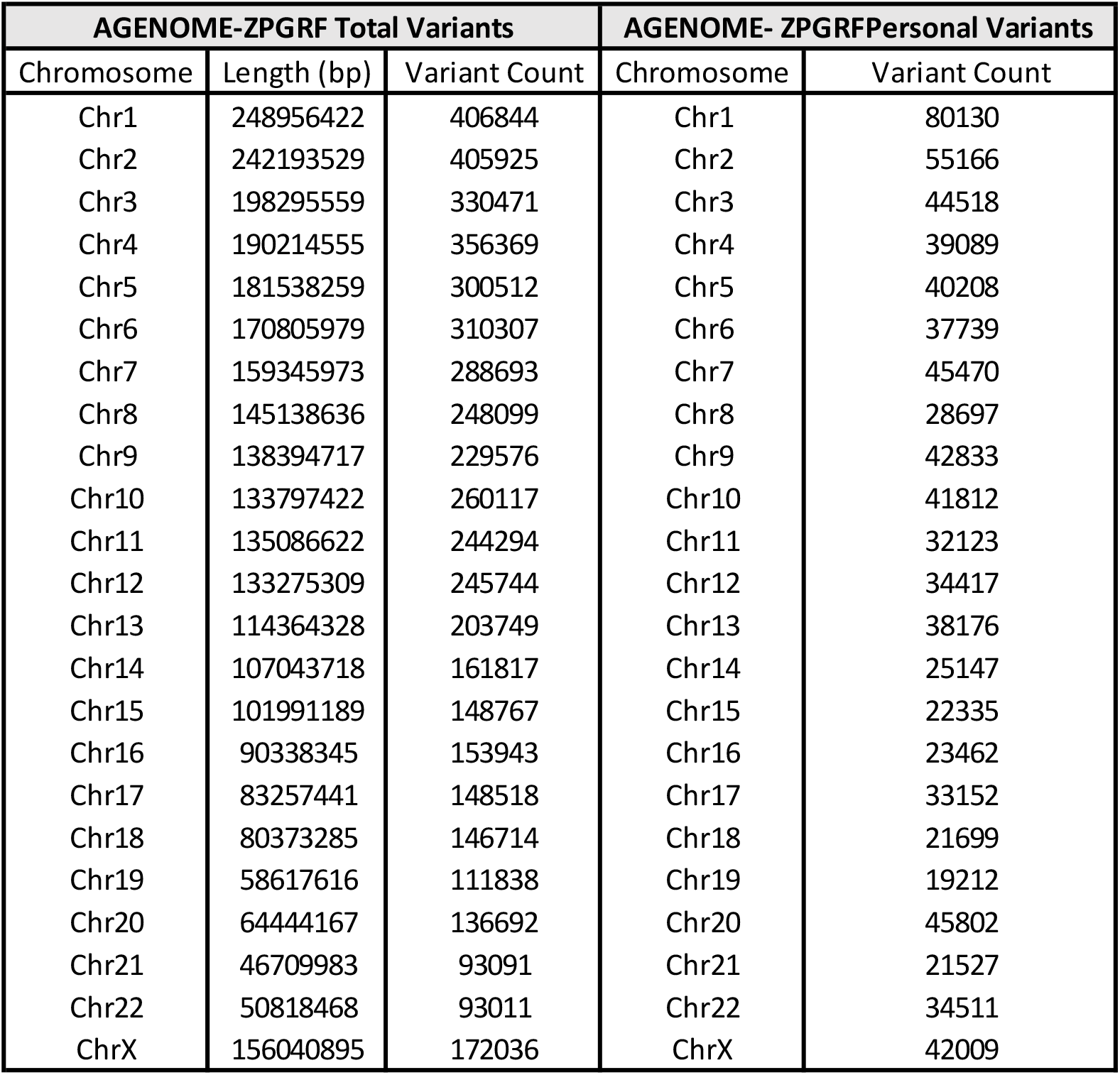
Chromosome-wise distribution of variants in AGENOME-ZPGRF and novel variants in AGENOME-ZPGRF

**Table 5:** Genomic location, functional class and impact of unique variants in AGENOME-ZPGRF

| Type (alphabetical order) | Count | Percent (%) |
| --- | --- | --- |
| DOWNSTREAM | 61074 | 7.176438102 |
| EXON | 4202 | 0.493751726 |
| INTERGENIC | 433381 | 50.92399255 |
| INTRON | 266354 | 31.29765521 |
| SPLICE_SITE_ACCEPTOR | 70 | 0.008225279 |
| SPLICE_SITE_DONOR | 51 | 0.005992703 |
| UPSTREAM | 78750 | 9.25343846 |
| UTR_3_PRIME | 5392 | 0.633581463 |
| UTR_5_PRIME | 1761 | 0.20692451 |
| TOTAL | 851035 |  |
| Number of effects by impact |  |  |
| Type (alphabetical order) | Count | Percent |
| HIGH | 415 | 0.043318195 |
| LOW | 1702 | 0.177656788 |
| MODERATE | 2150 | 0.224419562 |
| MODIFIER | 953760 | 99.55460545 |
| TOTAL | 958027 |  |
| Number of effects by functional class |  |  |
| Type (alphabetical order) | Count | Percent |
| MISSENSE | 2009 | 56.83168317 |
| NONSENSE | 31 | 0.876944837 |
| SILENT | 1495 | 42.29137199 |
| TOTAL | 3535 |  |

The high impact SNPs across the AGENOME-ZPGRF genome are distributed among 1,014 protein-coding genes in the genome and 311 non-coding regions. We next classified the high impact variants to understand the significance of the coding (sSNPs) and non-coding SNPs (nsSNPs), in terms of their distribution in protein class, pathways, biochemical activity with KEGG^43^ pathways using DAVID^44^. The distribution of both high impact sSNPs and nsSNPs was significantly enriched in G protein coupled receptor pathway genes, olfactory transduction. Our finding is consistent with studies that demonstrate higher levels of polymorphism observed in human olfactory gene family^45^. In addition, we found enrichment for pathways associated with neuroactive ligand-receptor and osteoclast differentiation. The majority of the high impact coding SNPs belonged to transmembrane helices, transmembrane and receptor protein classes. Reactome based pathway analysis^46^ showed that pathways for antigen presentation: Folding, assembly and peptide loading constituted the major pathway implicated for genes harboring the coding and non-coding SNPs **(Figure 4**).

### Genetic structural variation in AGENOME-ZPGRF

Using short-read sequencing data of AGENOME-ZPGRF, we called 69,148 SVs using DELLY2 structural variant prediction tool^33^ (**Figure 3**). We observed that while most of the SVs were deletions (n=40070), we found other SVs categorized as inversions (n=6004), duplications (n=5808), insertions (n=2129) and translocations (n=15137). No mis-assemblies were observed outside centromeric regions and segmental duplication regions.

**Figure 3:**
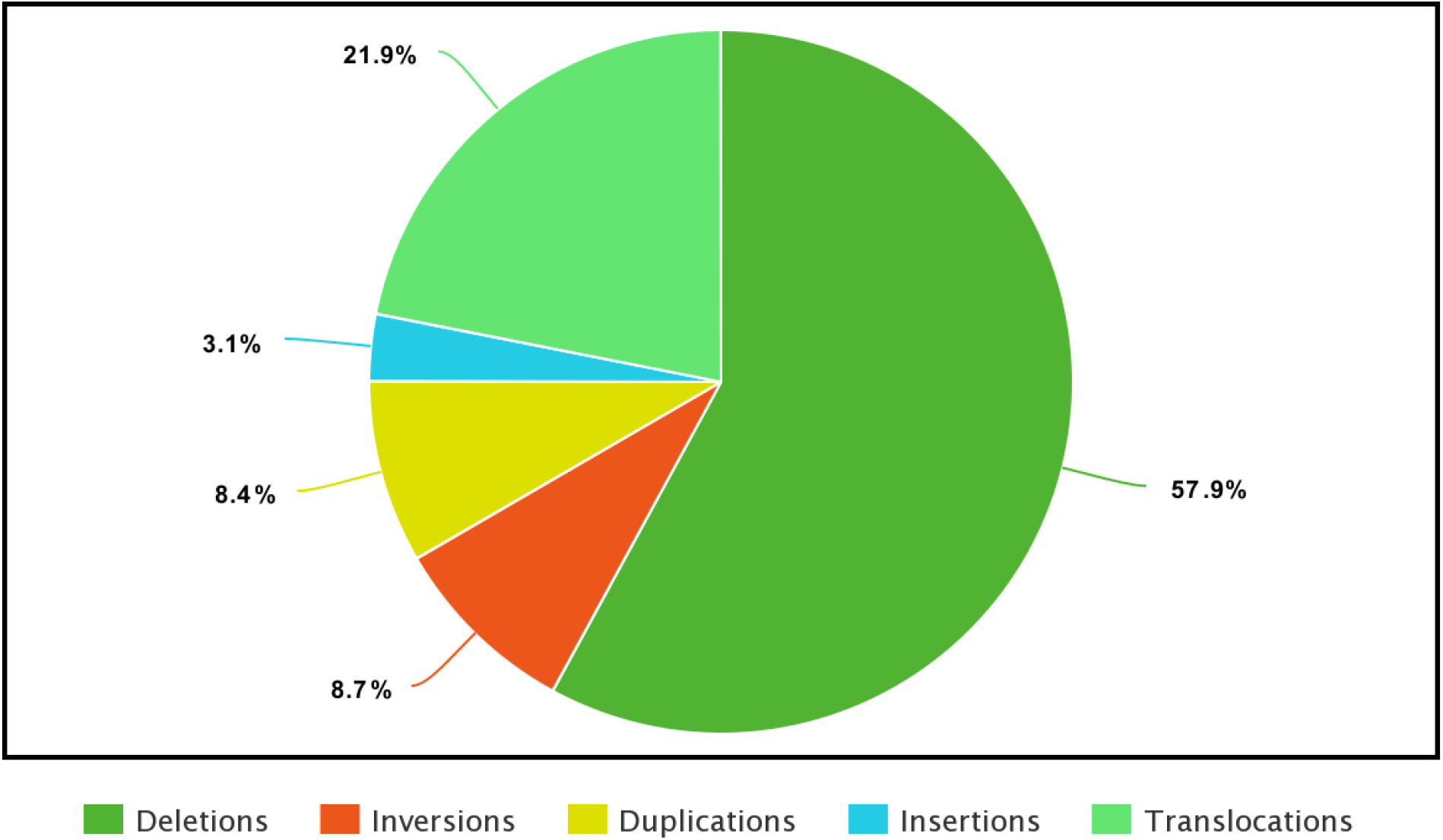
Distribution of Structural Variant (SV) calls in the assembled first Zoroastrian-Parsi Genome, AGENOME-ZPGRF; The breakdown of the SV’s is as follows: deletions (n=40070), inversions (n=6004), duplications (n=5808), Insertion (n=2129) and translocations (n=15137).

**Figure 4:**
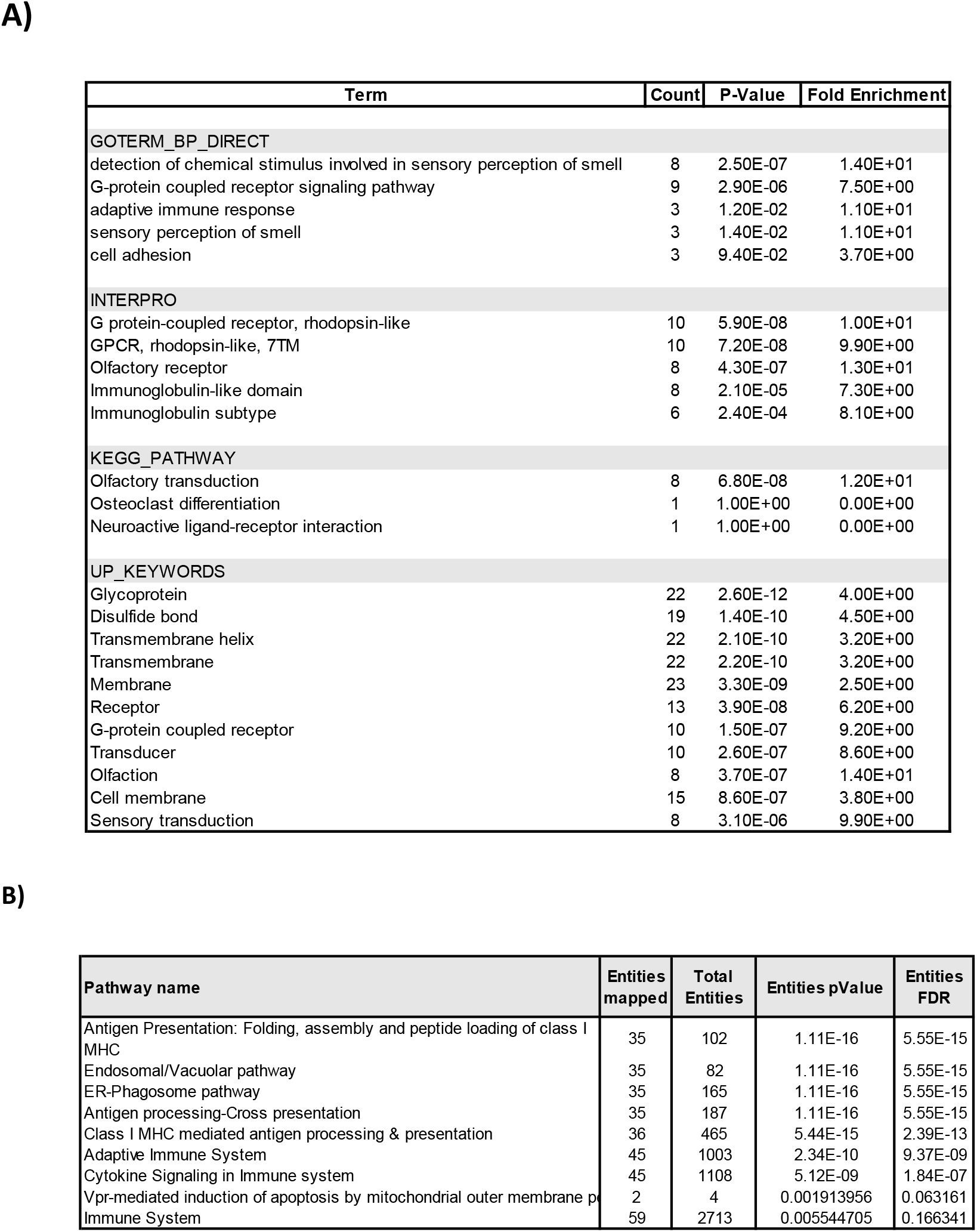
Enrichment of AGENOME-ZPGRF novel high impact variant across different databases like (A) DAVID and (B) Reactome

### Pharmacogenomics and drug risk assessment using AGENOME-ZPGRF SNPs

One of the aims of personalized genomics is to assess the individuals SNPs for disease and drug reaction assessment aiding drug dosage regimens. Using the variant-drug risk correlation annotation in the PharmGKB database^34^ and KEGG database, we sought to understand SNPs of pharmacogenomic relevance in the AGENOME-ZPGRF. We identified 20 unique SNPs (**Appendix 8**) associated with 12 genes distributed across 9 chromosomes (chr1,2,4,7,8,11,14,15 and 16) with pharmacogenomic relevance based on PharmGKB (**Appendix 7**). We identified 10 actionable SNPs from literature as it pertains to treatment with various drugs, some of which are also represented in the PharmGKB (Table 3).

## Discussion

We present the first, high depth whole Zoroastrian-Parsi Genome Reference Female sequence, AGENOME-ZPGRF for the Zoroastrian-Parsi population of India. The AGENOME-ZPGRF is a high depth whole genome sequence at 173X, combining genomic reads from short read (160X; Illumina) and long read (8X; 5X; Oxford Nanopore Technologies) sequencing technologies. AGENOME-ZPGRF represents the first high depth whole genome sequence from the Indian subcontinent, where there have been previous genome assemblies of male^47,22^ and female^21^ at <40X. AGENOME-ZPGRF, contains 2,778,216,114 nucleotides as compared to 3,096,649,726 in GRCh38. AGENOME-ZPGRF is unique as it is derived almost entirely from a single individual unlike GRCh38, which represents a mosaic of multiple individuals, thereby, adding further insight into personal genome and variant-disease association approaches. We have extended our study to sequence the first Zoroastrian-Parsi male genome, presently 2,589,561,354 bp in length with 87.46% genome fraction mapping to GRCh38 (**Appendix 9**).

The genome completeness is 93.25% which is on par compared to other high resolution whole genomes from the Ashkenazi^16^, Egypt ref^48^ and Yoruba genome^49^ assembly projects. Our assembly quality is further enhanced by BUSCO completeness score of 88.3% validating our benchmarking process for genome completeness. Annotation identified 20,833 genomic features, of which 14,996 are > 99% identical to their counterparts on GRCh38. Most of the remaining genes were partial. This assembly, designated AGENOME-ZPGRF, was the basis for all subsequent refinements. Our analysis revealed 5,426,310 variants of which 79% are SNPs (4,291,601) and 21% are indels, MNPs and mixed variants. AGENOME-ZPGRF had 960,867 novel/personal SNPs not listed on dbSNP. The AGENOME-ZPGRF reference standard of this endogamous, socially, genetically divergent community adds valuable insights into human population genetic diversity as compared to other global populations. Furthermore, our study adds information to the catalogue of genomic variation derived from the 1000 human genome project consortium, which also includes samples of Indian origin (1000 Genomes Project Consortium).

Besides the nuclear genome, we had previously studied AGENOME-ZPGRF mitochondrial genome variants^50^ as the first *de novo* Zoroastrian-Parsi mitochondrial genome, AGENOME-ZPMS-HV2a-1 (Genbank accession, <u>MT506314</u>). Our analysis showed that the AGENOME-ZPGRF belongs to haplogroup HV2a and that showed 28 unique variants compared with the revised Cambridge Reference Standard (rCRS). HV2a is an extremely rare haplogroup, and prevalent among the Zoroastrians-Parsis in our study cohort. The haplogroup HV2a is closely associated with Caucasian descent, with its documented prevalence dating back to ancient Scythians who were geographically distinct group of nomads joined by common cultural expressions^57^. They date back to about the 9th century BCE until the 4th century CE^56^ tracing their origins to the Caspian Pontic Steppes and the Altai mountains^51^, indicative of the unique genomic landscape of the contemporary Zoroastrian-Parsi among Indian and European communities.

Variants in personal genomes can be used to assess disease risk, carrier status and drug response/interaction contributing to a pharmacogenomic insights in clinical genetics. We have assessed the AGENOME-ZPGRF genome using OMIM, PharmaGKB and KEGG databases for SNPs with health and disease consequences. We identified high risk for Multiple sclerosis, among other diseases that include cancers and neurodegenerative diseases. We found two *CHRNA3, CHRNA5* alleles: rs16969968, rs1051730 gene variants that have been associated with cognition, possibly mediating in part risk for developing Nicotine Dependence^52,53^. We also found a C>A variant in *C11orf65* located near ATM gene regulating metformin response in Type 2 diabetics^54^. Additionally, we found intronic variant (rs762551) in *CYP1A2* associated with leflunomide induced toxicity in treatment for Rheumatoid Arthritis^55^. In the context of preventive pharmacogenomics association, we found the AGENOME-ZPGRF, harbored a SNP (C>T) in *DPYD;DPYD-AS1* implicated in fatal consequences to 5-Fluorouracil (5-FU)-based treatments (4%-5%, early onset-severe to 0.3%, fatal) in patients with dihydropyrimidine dehydrogenase (DPD) deficiency.

In sum, our present study has delivered the first, complete, *de novo*, high depth genome assembly AGENOME-ZPGRF for Zoroastrian Parsi community of India. Analysis of the variants in AGENOME-ZPGRF indicated 960,867 novel/personal SNPs, some of which were found associated with adverse drug interactions in separate studies. We have also completed the whole genome assembly of a Zoroastrian-Parsi male, whose variant annotation is underway. Further analysis of personal variant-disease-drug response annotations that are gender specific made using clinically validated variants will complement current healthcare practices with personalized pharmacogenomics. This will lead to safe, accurate, drug dosage and treatment regimen by physicians and clinical trials.

## Declarations

### Ethics approval and consent to participate

We would like to state that this ethics review board is not affiliated with a commercial entity, and we confirm that the Ethics Review was sought from an independent ethics review board not affiliated with the funder or the commercial entity, in line with Declaration of Helsinski that the “committee must be transparent in its functioning, must be independent of the researcher, the sponsor and any other undue influence and must be duly qualified”.

The study was approved by the Institutional BioEthics Committee constituted by the Department of Biotechnology, Government of India (BIAG-CSP-033). The committee constituted is compliant with the scientific, medical, ethical, legal and social requirements of the research proposal and in line with the 1964 Helsinki declaration and its later amendments. All subjects have provided written informed consent for the collection of samples and subsequent analysis.

### Competing interests

The authors declare that they have no known competing financial interests or personal relationships that could have appeared to influence the work reported in this paper.

### Funding

The project was funded by the grant awarded to Dr.Villoo Morawala-Patell, titled “Cancer risk in smoking subjects assessed by next-generation sequencing profile of circulating free DNA and RNA” (GG-0005) by the Foundation for a Smoke-Free World, New York, USA.

## Acknowledgements

We would like to thank Dr.Raja Mugasimangalam and Dr. Sudha Rao, Genotypic Technologies, for their valuable inputs, Dr.Paul Morill and Dr.Farah Patell Socha for their excellent project management and guidance regarding GG-0005, Dr.Sanaya Patell McGaw for editorial assistance and Ms.Mahima Kishinani, Ms.Janet Rymound for contributing to literature review and analysis. We thank the Zoroastrian-Parsi community of India for their enthusiastic cooperation and the The Avestagenome Project^®^ project team.

## Supplementary Figures

**Supplementary Figure 1:**
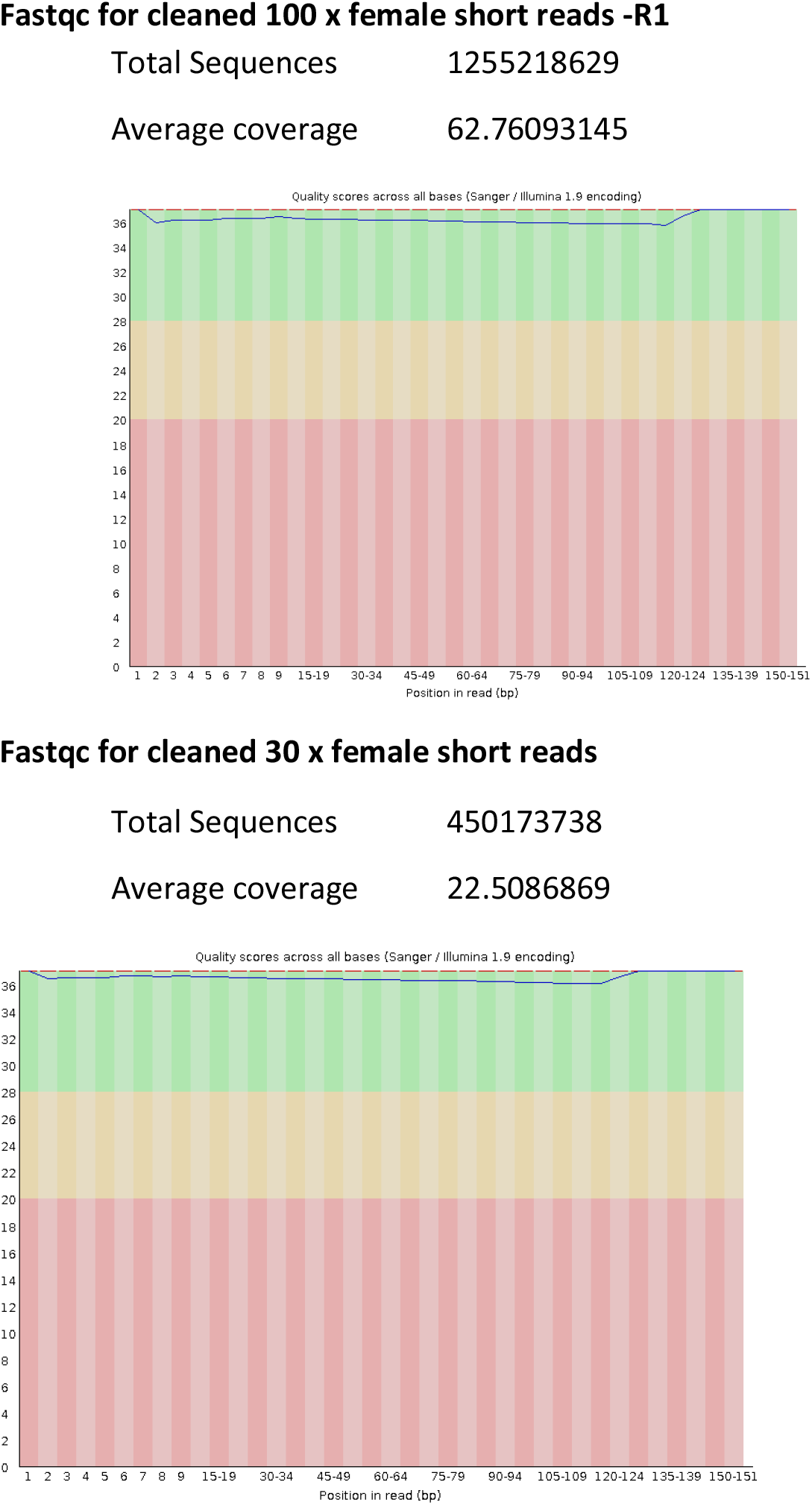
Representative read quality plot from FastQC after cleaning of reads using AdapterRemoval

**Supplementary Figure 2:**
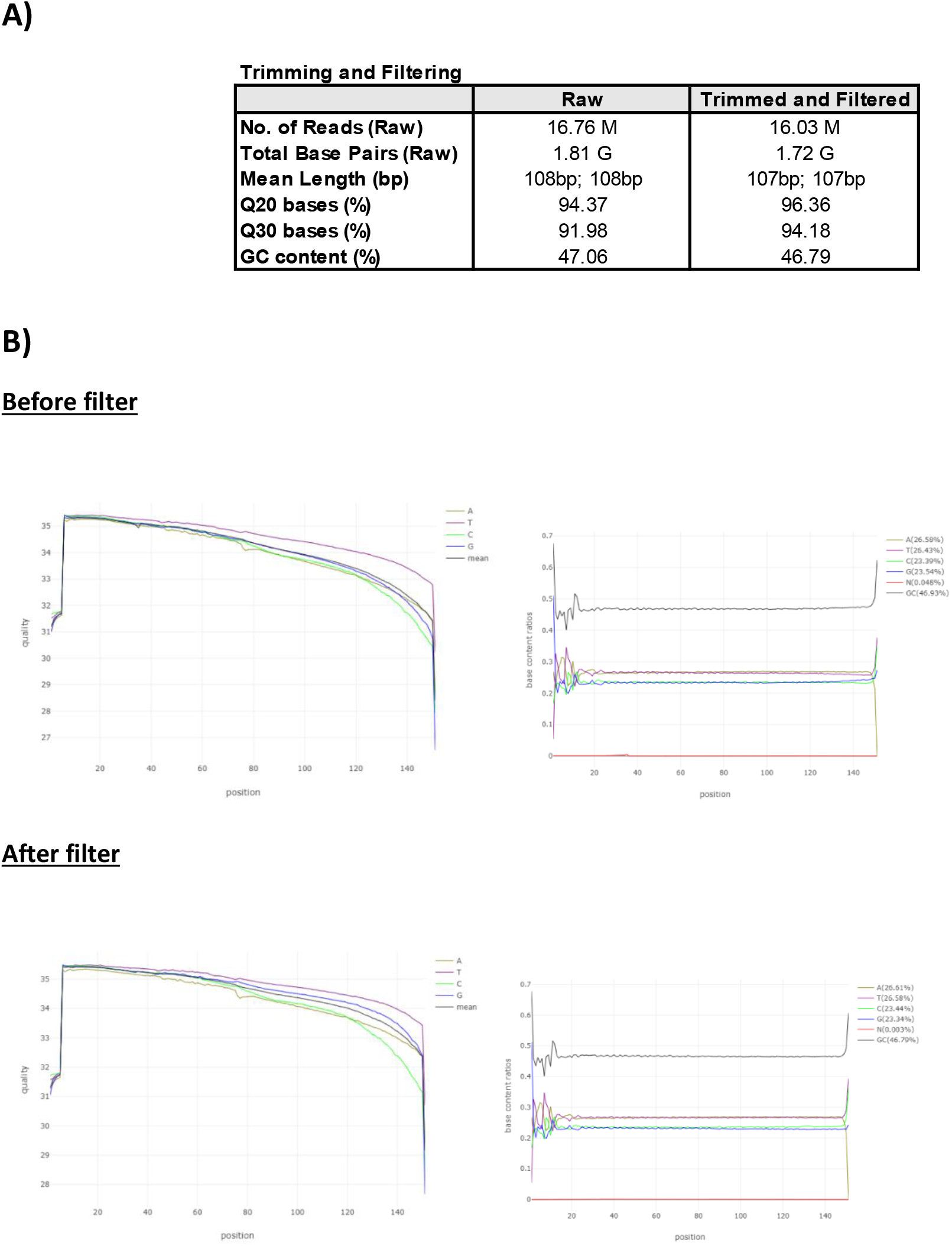
Read sequencing and Analysis statistics. A) Table indicating read processing and QC of sequencing data pre and post filtering B) Fastp output of read quality and base counts

**Supplementary Figure 3:**
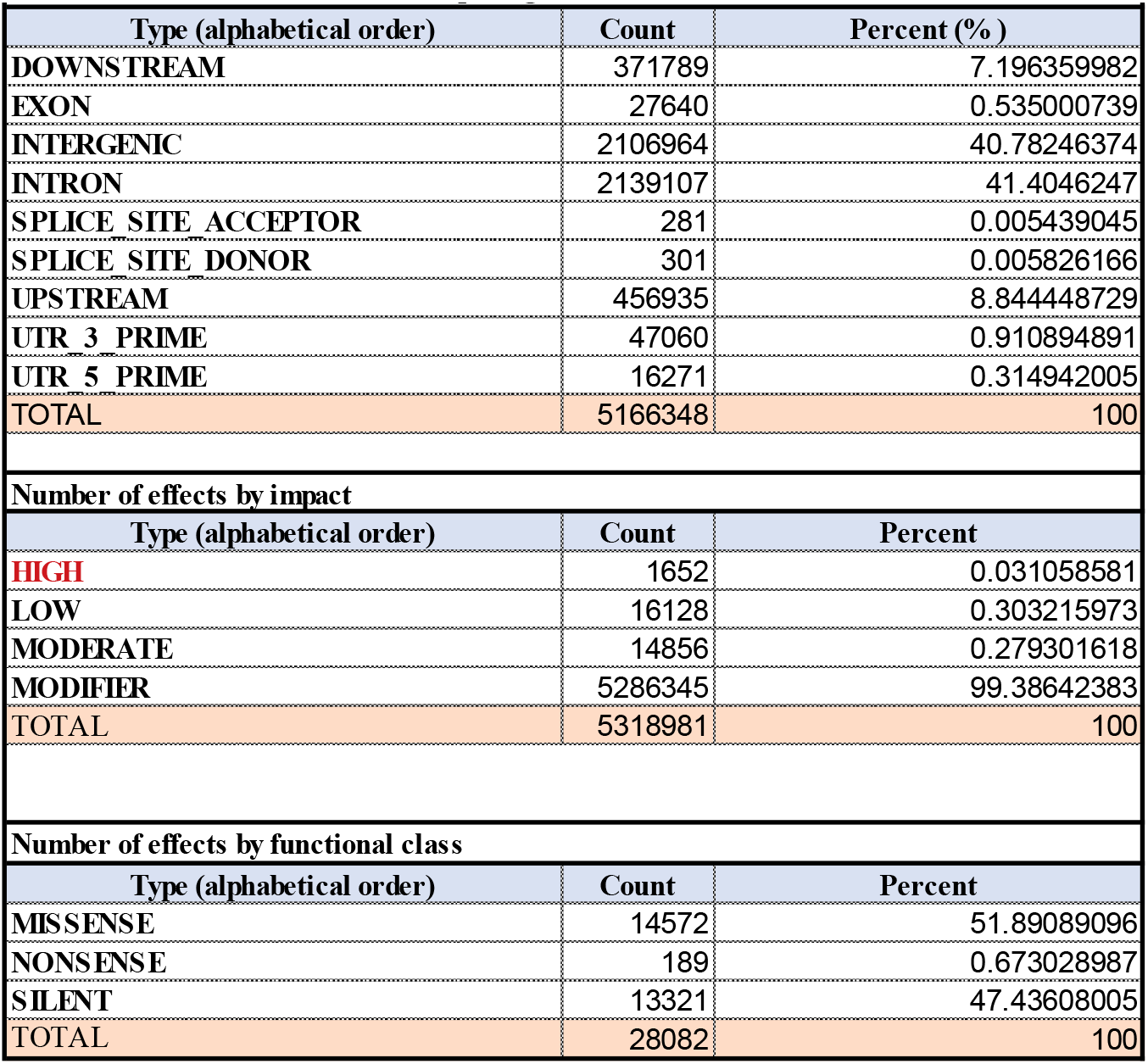
AGENOME-ZPGRF **g**enome wide distribution of variants across different genomic regions, by impact and functional classes

## Notes

### Competing Interest Statement

The authors have declared no competing interest.

